# An Associative Memory Hamiltonian Model for DNA and Nucleosomes

**DOI:** 10.1101/2022.10.15.512163

**Authors:** Weiqi Lu, José N. Onuchic, Michele Di Pierro

## Abstract

A model for DNA and nucleosomes is introduced with the goal of studying chromosomes from a single base level all the way to higher-order chromatin structures. This model, dubbed the Widely Editable Chromatin Model (WEChroM), is able to reproduce the complex mechanics of the double helix including its bending persistence length and twisting persistence length, and their respective temperature dependence. The WEChroM Hamiltonian is composed of chain connectivity, steric interactions, and associative memory terms representing all remaining interactions leading to the structure, dynamics, and mechanical characteristics of the B-DNA. Several applications of this model are discussed to demonstrate its applicability. WEChroM is used to investigate the behavior of circular DNA in the presence of positive and negative supercoiling. We show that it recapitulates the formation of plectonemes and of structural defects that relax mechanical stress. The model spontaneously manifests an asymmetric behavior with respect to positive or negative supercoiling, similarly to what was previously observed in experiments. Additionally, we show that the associative memory Hamiltonian is also capable of reproducing the free energy of partial DNA unwrapping from nucleosomes. WEChroM can readily emulate the continuously variable mechanical properties of the 10nm fiber and, by virtue of its simplicity, allows the simulation of molecular systems large enough to study the structural ensembles of genes. WEChroM is implemented in the OpenMM simulation toolkits and is freely available for public use.

**Author Summary:** The structural ensembles of genes have been so far out of the reach of theoretical and computational investigations because genes are molecular complexes too big to be tackled with even the most efficient computational chemistry approaches and yet too strongly affected by heterogeneous molecular factors to be effectively modeled as a simple polymer. In this work, we develop a computationally efficient, easy-to-use, and widely editable chromatin model to study the principles of DNA folding at the gene scale. Using the framework of Associative Memory Hamiltonians, this model reproduces the structural and mechanical properties of double-stranded DNA and accounts for the effects of nucleosome-forming histone octamers and other proteins bound to DNA. Our results open the path to studying the structural and mechanical ensembles of genetic systems as large as tens of kilobases of chromatin, i.e., the size of mammalian genes.

## Introduction

Deoxyribonucleic acid (DNA) has been studied for decades since the fundamental discovery made by Watson, Crick, Wilkins, and Franklin(1–3) The repeating units of the DNA molecule, nucleotides, are composed of phosphate, sugar, and a base group; a sequence of covalently bonded nucleotides form single-stranded DNA. The double-helix structure of DNA, formed by two strands of DNA, is maintained by base pairing, planar base stack interactions, and electrostatic interactions(4). In eukaryotes, DNA is then wrapped around histones to form nucleosomes, which serve as the basic structural unit of DNA packaging. Each nucleosome contains about 147 base pairs of DNA wrapped 1.7 times around a histone octamer. Recent evidence shows that the structure of nucleosomes is very dynamic and irregular(5). Outer stretches of nucleosomal DNA are observed to spontaneously unwrap and rewrap to the core octamer(6), which can result in DNA strands entering and exiting the nucleosome at flexible angles, and this ultimately has a significant impact on the three-dimensional packing of chromatin. However, we still lack a sufficient understanding of DNA and nucleosome mechanics to ascertain how these features combine with other factors to determine the structural, dynamical, and mechanical properties of the chromatin fiber and higher-order structures.

Numerous theoretical and computational approaches have already been explored to model DNA and chromatin fibers, greatly improving our understanding yet leaving many questions open. All-atom (AA) simulations have been extensively used to gain insights into DNA dynamics(7), flexibility(8), and other properties at a fine molecular scale. Fully atomistic models, however, cannot simulate DNA molecules at the level of nucleosome organization or chromatin fiber due to computational limitations, at least in the foreseeable future. Additionally, for very large molecular assemblies, such as even a short segment of chromatinized DNA, the accuracy of AA force fields remains an issue. This is particularly problematic in systems like DNA and histones in which ions play a determinant role. Coarse-grained (CG) approaches reach the time and length scales that are inaccessible to AA simulations. Many CG DNA models have been developed in recent years(9–22) following bottom-up or top-down procedures(4,23). In bottom-up approaches(9–13,24), effective interactions are determined to eliminate some degrees of freedom from reference atomistic simulations. As a result, CG force fields’ ability to reproduce the system’s behavior depends on the accuracy of the underlying AA force field, from which it inherits limitations in predictive capabilities, although reducing the computational cost. The top-down CG empirical energy functions(14–22) are tuned to match the properties of the system as observed through experiments, such as the polymer’s flexibility or melting temperature, thus avoiding the pitfalls of atomistic simulations altogether at the expense of missing some physical understanding. Examples of successful top-down CG models use a variable number of sites to represent a single nucleotide: from one(14,15), three(16–19), up to eight(21,22). The oxDNA model(14,15) represents each nucleotide as a rigid body with 3 interacting sites—one site for each pair of phosphate and sugar groups and two sites for each base. The effective potential is fitted to reproduce the experimentally determined structure and mechanics of single- and double-stranded DNA as well as the thermodynamics of hybridization. The 3-site-per-nucleotide (3SPN.0(16)/1(17)/2(18)/2C(19)) model was developed by de Pablo’s group to reproduce experimental persistence length for both single-stranded DNA (ssDNA) and double-stranded DNA (dsDNA), DNA melting temperatures, and hybridization rate constants. The most recent version, 3SPN.2C(19), also incorporates sequence-dependent shape and mechanical properties by introducing new weak dihedral potentials.

Coarser models are necessary to perform simulations of the chromatin fiber architecture. A mesoscale model developed by Schlick and coworkers(25,26) has been used to investigate the relationship between chromatin organization and structural factors, such as nucleosome repeat length and linker histones(27–31). The model describes the nucleosome core particle and its charge distribution, the histone tails and the linker DNA that is modeled using a single particle for every 10 bp. Lequieu et al.(32) developed a coarse-grained model, 1CPN, where histones, including their tails and the DNA wrapped around them, are represented as one ellipsoid. Each 3 bp of linker DNA is described by a particle with orientation (i.e., 6 degrees of freedom) in a twistable wormlike chain. The model is tuned to approximate the free energies and dynamics of single nucleosomes and short chromatin fibers.

Traditional CG models involve two types of interactions: bonded potentials among neighbors and non-bonded potentials among any pair of particles. The bonded interactions typically entail the 2-body bonding term, 3-body angle or bending term, 4-body aligning/twisting/dihedral terms, and 4-body to many-body terms to capture base stacking effects. Non-bonded interactions include excluded-volume potentials and electrostatic potentials where the solvent is typically treated implicitly. While following a top-down philosophy, traditional CG models aim to reproduce the physicochemical nature of DNA using interaction potentials motivated by basic chemistry; however, such ambitious effort is often frustrated by a very difficult parametrization process and computationally expensive simulations, due to multi-body interactions and non-bonded electrostatic potentials.

Such detail may not be necessary if we limit our objective to recapitulating the mechanical properties of DNA. Here, we use an Associative Memory Hamiltonian (33) (AMH) to explore the energy landscape of chromatinized DNA, starting from known structures of DNA fragments and protein-DNA complexes. The AMH framework has been successfully employed in protein folding(33) where the energy landscape of a given protein is reconstructed using crystallographic information of short fragments characterized by sequences overlapping that of the protein under investigation(33). We apply this data-driven approach to chromatin folding, building a DNA Hamiltonian from the computational structure of B-DNA and crystallographic structures of nucleosomes. The AMH implicitly recapitulates all the energy terms already discussed: bending, twisting, base-stacking, and electrostatic energy. Parameters are tuned to reproduce the mechanical properties of DNA, including the twisting and the bending persistence lengths, and their coupling with supercoiling behaviors.

This model, dubbed the Widely Editable Chromatin Model (WEChroM), represents each nucleotide as a single particle and, besides polymer connectivity, uses AMH energy terms for all bonding interactions. By virtue of its simplicity, the WEChroM is computationally efficient, and it allows simulating the dynamics of large genetic molecular systems up to kilobases of chromatin at base-pair resolution, reaching approximatively the size of mammalian genes. Therefore it will also allow connections to large scales chromatin structures, including those observed experimentally by chromosome capture methods(34–37) and imaging techniques (38–41), and those simulated using phenomenological polymer models(42–44).

We show that the WEChroM accurately reproduces the properties of naked DNA properties, such as bending and twisting persistence lengths, as well as supercoiling behaviors. Because of the finite cost of the AMH potentials, the model can also account for the natural emergence of defects in the structure of DNA. Lastly, by introducing an associative memory template for the histone octamer, we also show that WEChroM recapitulates the free energy profile of nucleosome unwrapping.

## Results

### An Associative Memory Hamiltonian for DNA

The AMH was introduced by Friedrichs and Wolynes(45) to study the problem of protein folding. Using crystallographic information, the AMH reconstructs the energy landscape of a protein and encodes the native structure of the target sequence as the global attractor of such a landscape. The energy landscape of the protein is approximated by selecting a subset of the pairs of atoms in the reference protein structure and constraining their distances using a Gaussian well potential. The resulting model of the molecule is roughly harmonic in the folded state but can accommodate configurations with partial or complete unfolding of the protein at a finite energy cost. More than one molecular configuration can be encoded in the Hamiltonian; each one of these configurations is called a “memory”, in analogy with the terminology used in the field of recurrent neural networks(33,45,46).

We adapt the AMH framework to reconstruct DNA interactions and DNA-protein interactions. We start from idealized computational structures for B-DNA and from crystal structures of DNA-protein complexes as memories to construct the energy landscape of DNA and nucleosomes. In the WEChroM, each nucleotide is represented by one particle, as shown in Fig 1. The model is sequence-independent, but this can be easily generalized. The resolution of one nucleotide per particle allows for describing both the bending and twisting properties of DNA without additional rotational coordinates, and it allows the model to closely reproduce the complex mechanical behavior of the double helix, including buckling and the formation of plectonemes.

**Figure 1.**
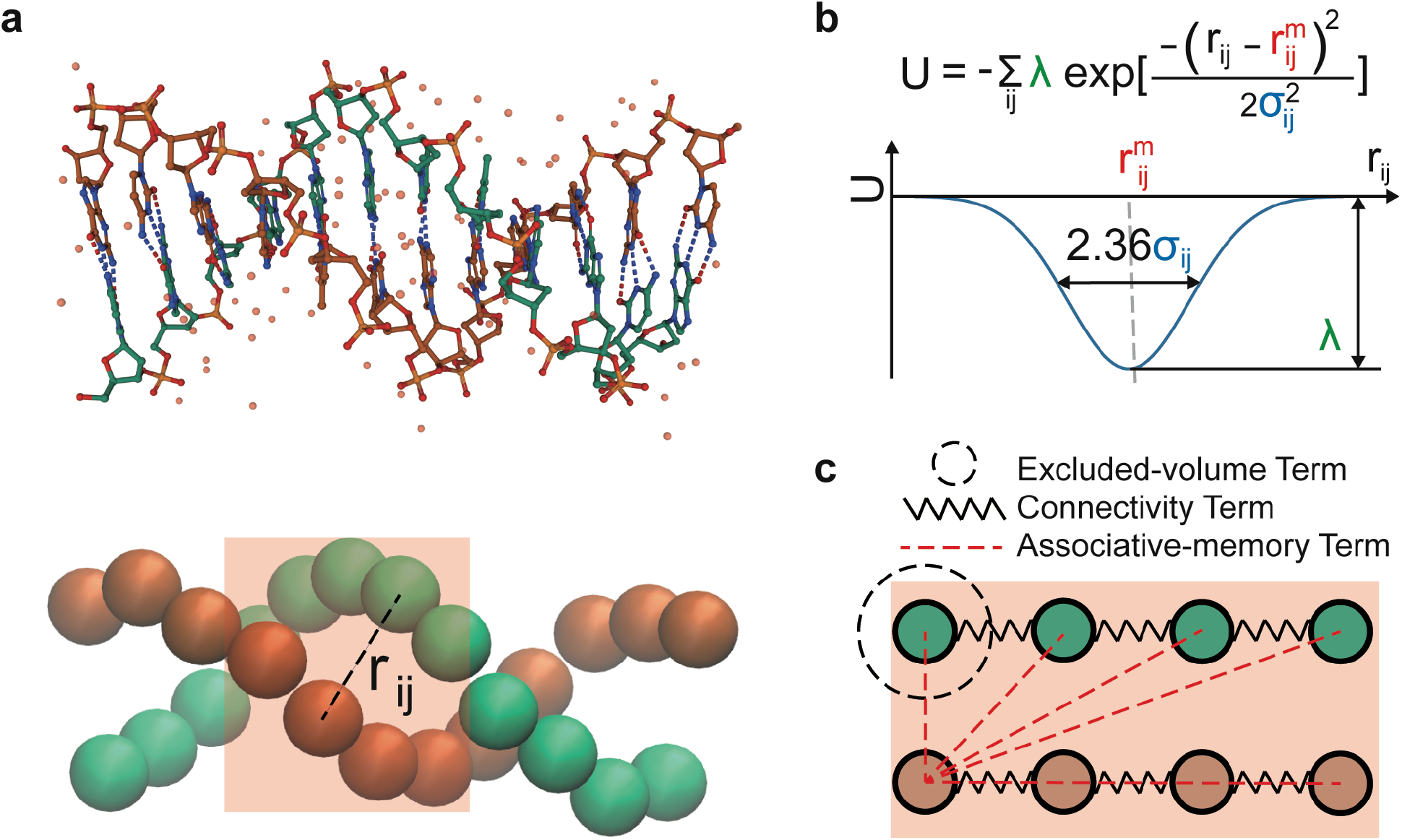
An Associative Memory Hamiltonian for DNA. (a) Comparison of an atomistic representation and a coarse-grained representation of a B-DNA dodecamer (PDB ID: 1BNA). (b) The AMH recapitulates the energy landscape of the DNA molecule using a series of Gaussian wells. The parameters 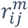 are determined using the distances measured in the structural memories, while the energy of interactions, λ and the scale of structural fluctuations, σ, are tuned to reproduce experimental observables. (c) Schematic representation of the energy terms in the model: connectivity (springs), excluded-volume terms (dashed circle), and AMH terms encoding the double helix structure (red dashed lines). An AMH term is applied between any pair of particles inside the box of 4bp. The box is then slid along the DNA, and the procedure is repeated.

For naked DNA, the WEChroM potential energy consists of 3 major components: chain connectivity, steric interactions, and associative memory interactions, in this case labeled the double helix term (*U*_*DH*_). The chain connectivity is enforced by harmonic interactions between two neighboring particles on the same strand, mimicking the covalent backbone bonds in a single-stranded DNA. Steric interactions are modeled using a short-range repulsive potential. A complete description of the energy terms modeling chain connectivity and steric interactions is provided in the S2 Appendix.

Pairwise distances 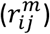 among nucleotides are captured from computational B-DNA structures and define the associative memory energy term (*U*_*DH*_)

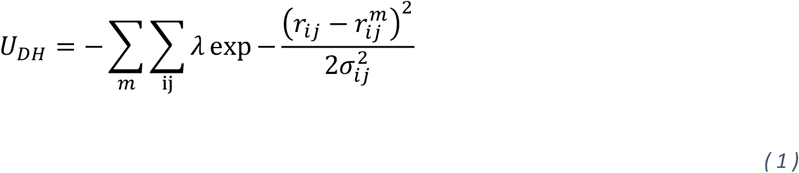

where *r*_*ij*_ is the distance between nucleotides *i* and *j*, and where the outer sum runs over memories and the inner sum over all pairs of nucleotides within the memory. The energy *λ* determines the strength of interactions and it is chosen to be a constant value, regardless of the nucleotide identity. The variance *σ*_*ij*_ controls the scale of structural fluctuations. To allow for more flexibility when the interaction distance is larger, *σ*_*ij*_ increases proportionally to the square root of the genomic distance between *i, j*, i.e., 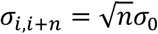, (Fig. S1).

In our model, each memory comprises 4 consecutive bp (Fig. 1c). Our analysis indicates that 4bp is the minimal memory size needed for reproducing the mechanics of DNA without creating artifacts. While we were able to reproduce the bending and twisting persistence length of DNA using 3bp memories, this required unreasonably large interaction energies between nucleotide pairs, and we thus rejected the model. We introduce energy terms by sliding the 4-bp memory box along DNA in 1-bp shifts. This results in multiple memory interactions being added between pairs of nucleotides depending on their index difference (Fig. S1), with pairs of nucleotides closer in genomic distance interacting more strongly.

To create a sequence-independent structural template, we choose 10 random sequences and generate their structures using X3DNA(47); we then calculate each nucleotide’s center of mass and use it to define the distances between nuclei 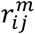. *U*_*DH*_ enshrines the double-helical structure of DNA as the global energy minimum of the WEChroM Hamiltonian while allowing for structural distortions as well as local defects in the DNA polymer, such as kinks and bubbles.

The current model does not include any energy term enforcing chirality. The correct chirality is set by the initial condition and preserved throughout the motion. While in principle energy fluctuations could change the global chirality of the system, in practice we did not observe any of these events during the extensive testing of the model. We concluded that a term enforcing chirality was not necessary, at least for the systems under investigation. It is possible, however, that future studies for particular larger molecular systems and different conditions might require such an energy term.

Similar to modeling double-stranded DNA, we also use the associative memory Hamiltonian framework to describe how proteins influence the structure of chromatin. We model the effect of the histone octamer on DNA by using the crystal structure of a nucleosome core particle (PDB ID: 1kx5) as the associative memory template. In this case, associative memory terms are introduced among nucleotides and between them and the histone octamer’s center of mass. Despite the simplified representation of protein-DNA interactions, our model recapitulates the free energy changes involved in the partial unwrapping of DNA from nucleosomes.

In summary, the AMH is a flexible theoretical framework that allows for introducing structural memories from a vast repertoire of DNA-binding proteins and histone variants and integrating the effect of these molecules at larger scales. This namesake characteristic of the Widely Editable Chromatin Model makes it particularly suitable for studying genetic systems above the nucleosomal scale and below the chromosomal scale, bridging the gap between the atomistic and bottom-up models typically utilized up to the nucleosomal scale, and the polymer physics approaches typical of the chromosomal scale(42). The WEChroM is thus designed to provide the means for studying the structural ensembles of genes, which are too large for conventional molecular modeling and yet strongly affected by heterogeneous molecular factors to be effectively modeled by simple polymer models.

### Bending and Twisting Persistence Lengths

The molecular rigidity of naked DNA is quantified by two persistence lengths: the bending persistence length (*L*_*bp*_) and the twisting persistence length (*L*_*tp*_). Experimentally, the bending persistence length *L*_*bp*_ of DNA has been found to be approximately 150 bp (around 50 nm) (50), while the twisting persistence length *L*_*tp*_ is believed to be between 75 and 360 bp (or around 25—120 nm)(51–53). In our model, the bending persistence length *L*_*bp*_ is defined by the relaxation of the angle correlation as a function of the genomic distance L:

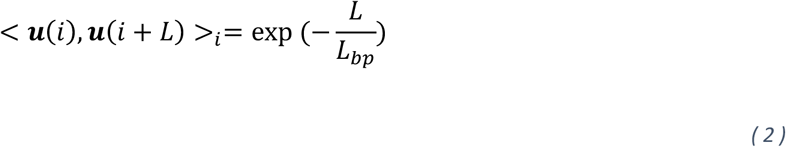

where ***u***(*i*) is tangent to the helical axis at base-pair *i*. Due to DNA’s double helix structure, the angle formed by any two base pairs is oscillating, leading to the following relaxation function for the twisting angle:

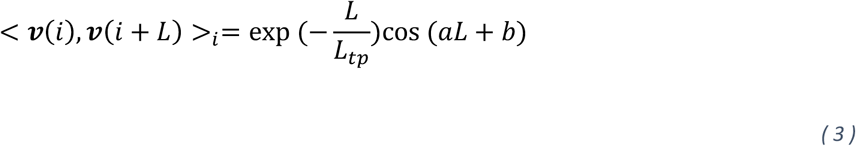

where ***ν***(*i*) is the vector pointing from one strand to the other strand within base pair (*i*). The twisting persistence length *L*_*tp*._ is defined from the exponentially decaying envelope 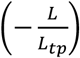 while *a, b* are the period and phase of the double helix.

By tuning the parameters *λ* and *σ*, in the double helix term in the AMH, we obtain a range of values for *L*_*bp*_ and *L*_*tp*_ as shown in Fig. 2a and Fig. 2b, respectively. The bending persistence length *L*_*bp*_, observed in simulations varies between 100 to 500 bp and monotonically increases with increasing *λ* or decreasing *σ*. This is expected as larger *λ*′*s* lead to stronger associative memory interactions while smaller *σ*′*s* lead to reduced fluctuations of the distances between nucleotides, both resulting in a stiffer molecule. The twisting persistence length *L*_*tp*_, observed in simulations varies between 89 and 415bp, in good agreement with the experimentally determined range between 75 and 360bp (Fig. 2b and Fig. S3). The persistence lengths obtained simulating systems of different sizes (Fig. S3) are stable, indicating the convergence in the simulations.

**Fig 2.**
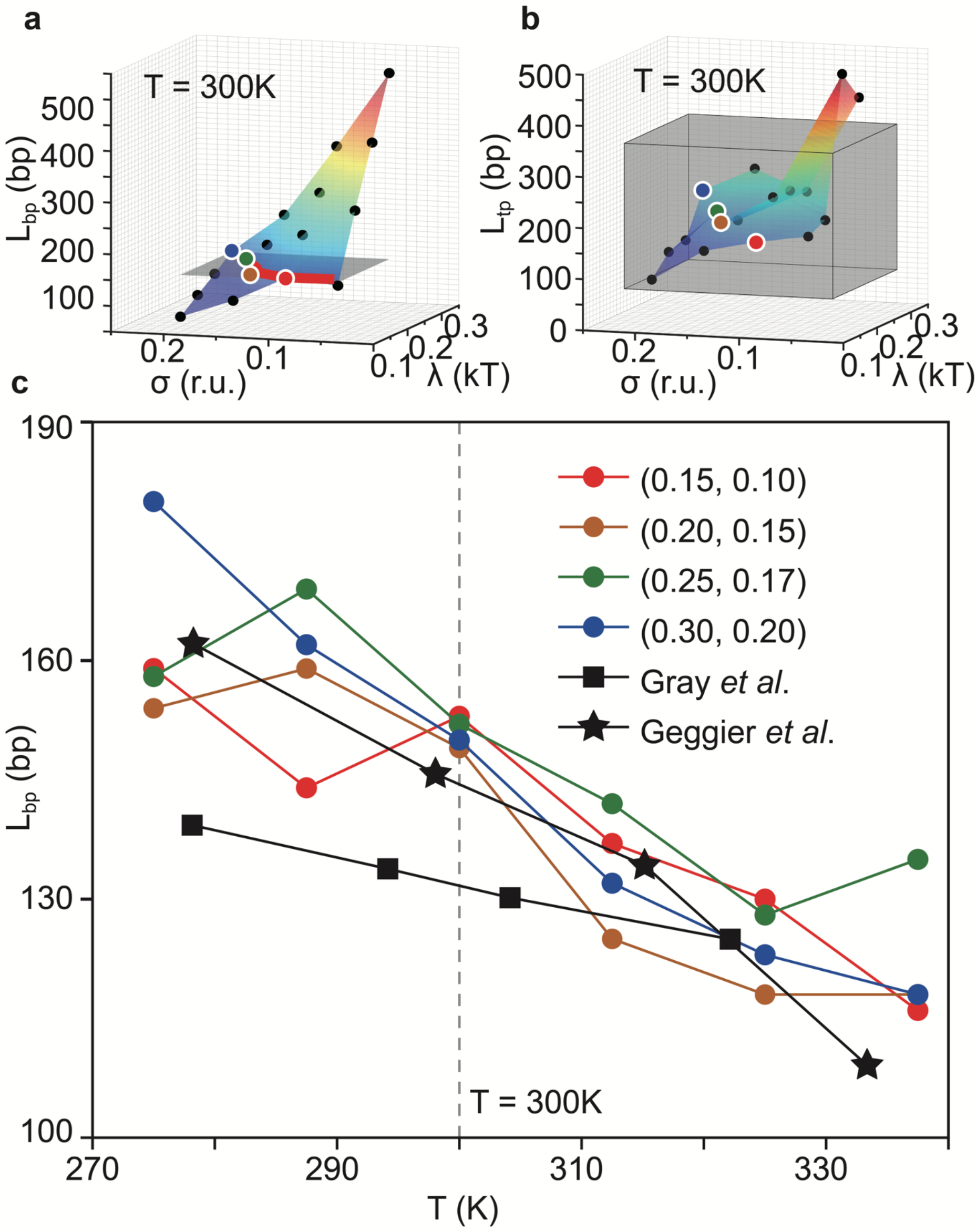
DNA Persistence Length and Parametrization. (a)(b) These figures show how persistence lengths depend on the strength of the energy interactions, λ, and the scale of structural fluctuations, σ. The bending persistence length is shown in panel (a), while the twisting persistence length is shown in panel (b). The σ is in reduced units (r.u.) which are explained in the S1 Table. Dots indicate the data points obtained by simulation; colored surface indicate their linear interpolation. The horizontal gray plane in (a) shows the experimentally determined bending persistence length of DNA, 150bp. This persistence length is set as the target for the optimization of the parameters. The red line indicates the intersection of the two surfaces, where our model meets the target persistence length value. Similarly, the gray box in (b) is the experimentally determined twisting persistence length, 75-360bp, our target in this case. (c) The figure shows the temperature dependence of bending persistence length. The four colored lines show results obtained using the WEChroM model. Black lines show experimental results published by Gray et al.(48) and Greggier et al. (49), respectively. As shown, the proposed model agrees well with the experimental observations for several couples of parameters (λ, σ).

Experimental studies have determined that the DNA persistence length depends on temperature, with *L*_*bp*_ ranging from 140^∼^165 bp at 277K to approximately 110^∼^120bp at 330K (48,49). We chose parameter sets characterized by a *L*_*bp*_ ^∼^150bp at 300K, i.e., consistent with experiments at room temperature. For these parameter sets, we calculate the persistence length *L*_*bp*_ as a function of the temperature, ranging from 270K to 340K. Simulations show that the persistence length *L*_*bp*_ decreases from 150^∼^180 bp at 275K to 120^∼^140bp at 338K, which is in excellent agreement with the experimentally determined temperature dependence (Figure 3c).

**Fig 3.**
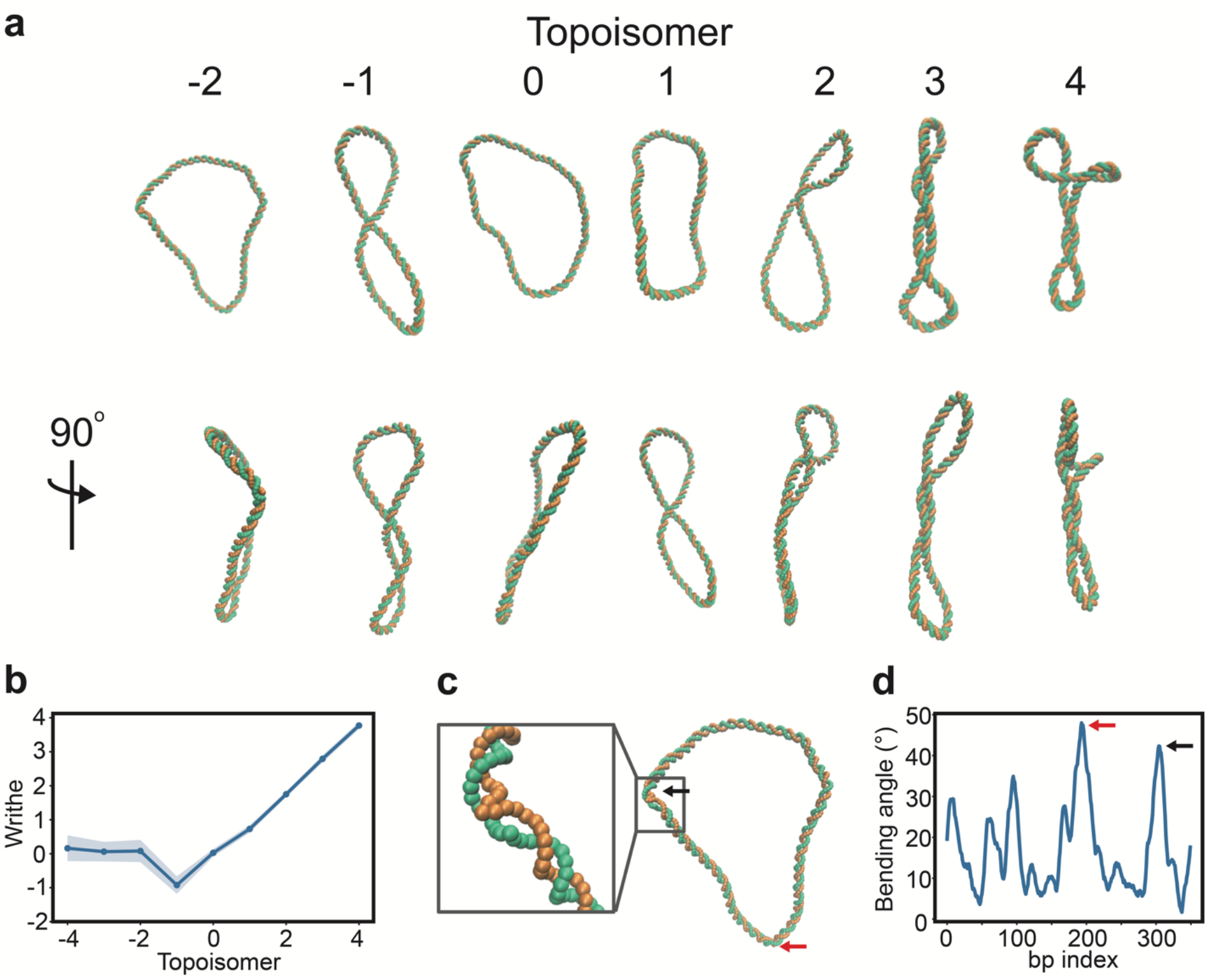
Supercoiling behavior of DNA minicircles. (a) Commonly observed shapes of DNA minicircles (b) Writhe analysis of DNA topoisomers -4 to 4. Positively supercoiled minicircles relax torsional stress forming plectonemes, which manifest in increasing writhe and are visible in the DNA conformations in (a). Moderately negatively supercoiled minicircles also form plectonemes, this time resulting in negative writhe. Strongly negatively supercoiled minicircles do not appear to form plectonemes. (c) Negatively supercoiled DNA relaxes torsional strain through local melting of the DNA double helix. The figure shows an example structure of a topoisomer -2, defects are indicated by the black and red arrows. (d) Bending angle along DNA for the minicircle structure in (c). Local defects are clearly visible at index 304 and 193, once again indicated by the black and red arrows.

Multiple sets of parameters are compatible with the reported bending and twisting persistence lengths from experiments (highlighted in Fig. 2a and Fig. 2b and listed in Table S2). These different sets of parameters, although having similar persistence lengths, show different features in other aspects. For example, the parameter set with (*λ, σ*) = (0.15, 0.05) has a similar *L*_*bp*_ and *L*_*tp*_ as other data points as shown in Fig. 2a, yet the double helix with this parameter set is more prone to forming defects at room temperature. Nonetheless, other parameter sets can produce *L*_*bp*_ and *L*_*tp*_ that are consistent with the experiments without obvious defects. Such degeneration is caused by the scarcity of experimental constraints and can be removed by introducing additional information on specific genetic systems.

We performed a sensitivity analysis of both bending and twisting persistence length with respect to the energy terms in our model (Fig. S2). Such an analysis revealed that the stiffness of the double helix is much more sensitive to the strength of the inter-strand interactions than to changes in the intra-strand interactions, with minor increases in the strength of inter-strand bonds leading to a significantly stiffer DNA polymer. This is consistent with the fact that the bending persistence length of a single-stranded DNA (ssDNA) is only approximately 6 bp(54), much smaller than that of double-stranded DNA (dsDNA).

### Supercoiling Behavior of DNA Minicircles

Genomic DNA is often subjected to torsional stress, which can both over- and under-wind the DNA double helix(55), resulting in twisting and coiling of the helix. Supercoiling can be induced by enzymes in order to reduce the volume of chromatin. Therefore, it will also be inevitably introduced when part of the DNA is uncoiled during essential cell processes, like transcription or replication(56,57). Elucidating the structural properties of DNA molecules in different supercoiling states is therefore essential to improve our understanding of genome functions.

In order to quantitatively characterize supercoiling, we utilize the linking number (*Lk*), i.e., the total number of times the two single DNA strands coil about one another. The linking number is related to two geometrical properties of the molecule, the twist (*Tw*, the coiling of the two strands about the helical axis) and the writhe (*Wr*, the coiling of the helix axis’s path around itself). These three properties are related by the simple relation

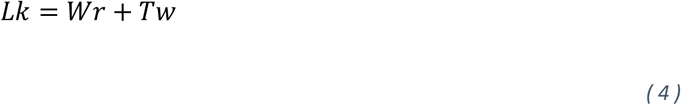

More details on how to calculate these quantities are found in S4 Appendix.

When the ends of a DNA molecule are covalently ligated to form a circle, the two strands become intertwined and will remain in such a state unless one of the strands is broken, i.e., the linking number is conserved. We performed simulations of supercoiled 350 bp DNA minicircles using the WEChroM model. We prepared 9 topoisomers with both negative and positive supercoiling, ranging from Δ*Lk* = −4 to Δ*Lk* = 4; the topoisomer Δ*Lk* = 0 corresponding to the relaxed DNA minicircle.

Similarly to what is seen in experiments (58), a wide variety of structures is observed in simulations (Fig. 3a). Positively or moderately negatively supercoiled minicircles, Δ*Lk* = −1 to Δ*Lk* = 4, tend to form plectonemes. These topoisomers transpose twist into writhe, relaxing twisting torsional strain into the bending strain due to writhe (Fig. 3b). Surprisingly, strongly negatively supercoiled DNA minicircles do not show a tendency to form plectonemes. On the contrary, negative torsional stress appears to be more easily relaxed through the formation of defects in the double-helical structure, i.e., by local DNA melting (Fig. 3c). Such asymmetric behavior of DNA with respect to supercoiling was previously observed experimentally (59,60), with AFM and Cyro-EM showing that negatively supercoiled minicircles often present melted regions at room temperature while positive ones do not (58,59,61).

As shown, the energy transfer between bending and twisting modes is an essential aspect of DNA mechanics. Such energy transfer is difficult to reproduce when using simpler models that lack details about the three-dimensional nature of DNA. Additionally, the formation of defects in the double-helical structure of DNA also appears to be essential to correctly describe its structural ensembles. Taken together, these two facts highlight the necessity of two crucial design elements of the WEChroM: accounting explicitly for the 3D structure of the DNA double helix and using a non-harmonic potential to recapitulate its empirical energy function.

### DNA-Protein Interactions and Nucleosomes

In eukaryotes, DNA is wrapped around histones to form nucleosomes with each nucleosome containing approximately 147 base pairs of DNA wrapped 1.7 times around a histone octamer. Besides compacting chromatin, nucleosomes create a significant barrier to protein binding, in turn contributing to the regulation of essential cellular processes like the gene expression (62). The outer stretches of nucleosomal DNA are observed to spontaneously unwrap and rewrap to the core octamer(6), significantly impacting the three-dimensional packing of chromatin. A faithful description of nucleosome dynamics is necessary to recapitulate the mechanical behavior of the chromatin fiber.

We introduce nucleosomes in the WEChroM by representing the histone octamer with a single particle at its center of mass, then apply AMH interactions between the DNA and histone particles to reproduce the effect of protein-DNA interactions. The DNA-protein interactions *U*_*Nucleosome*_ can be formulated as follows:

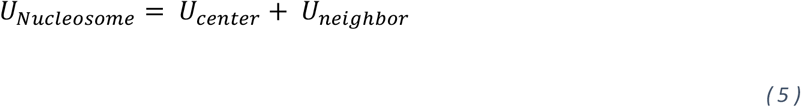

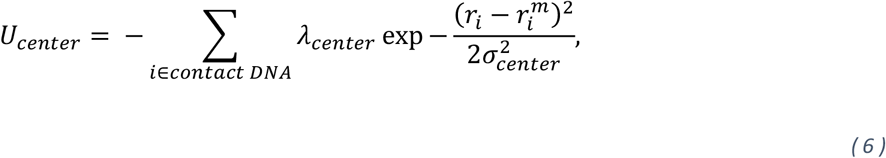

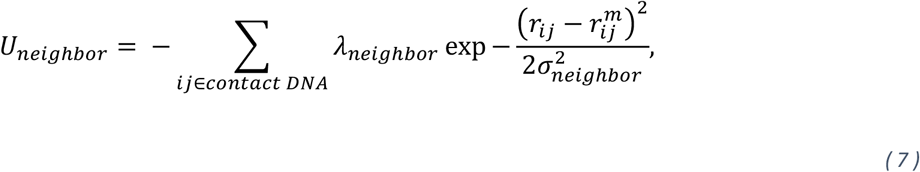

DNA bases in contact with the amino acids of the histone octamer are defined as “contact DNA”. The first part of the nucleosome associative memory *U*_*center*_ acts between the center of histone core and contact DNA (Figure 4a). The distances 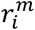 are learned from a nucleosome crystal structure. The second potential, *U*_*neighbor*_, acts between neighboring contact DNA bases. The energy *λ* and the variance *σ* in the two parts were parameterized to match the thermodynamics of the isolated nucleosomes.

**Fig 4.**
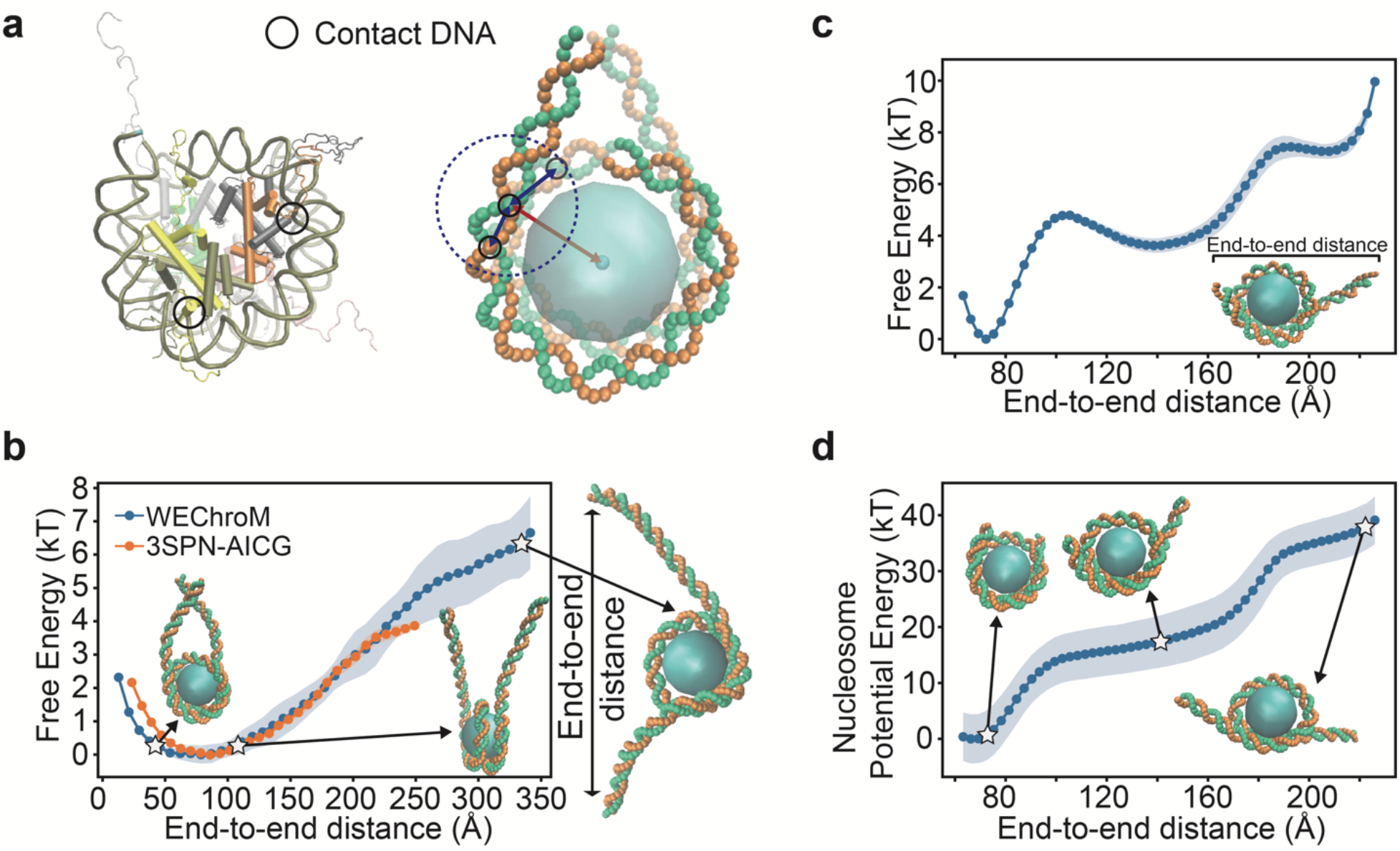
Nucleosome model and unwrapping analysis. (a) A detailed representation of the nucleosome core particle (PDB ID: 1KX5) (left) and the corresponding coarse-grained WEChroM representation (right). In the left figure, the upper half of the nucleosome particle is shown as solid while the lower half is transparent for clear visualization. The DNA nucleotides in close contact with amino acids are defined as “contact DNA” and subject to the AMH potential modeling histones-to-DNA interactions (black circle). In our model, contact DNA interacts with the histone octamer’s center of mass (U_center_, represented by the red arrow in the figure). Contact DNA also interacts with neighboring contact DNA (U_neighbor_, represented by the black arrows inside the indigo dashed circle). (b) Free energy of partial DNA unwrapping for a system composed of the histone octamer plus 223-bp of DNA; WEChroM model (blue) and 3SPN-AICG model (orange) Example nucleosome configurations are shown for end-to-end distances of roughly 45 Å (left), 107 Å (right), 330 Å (outside) (c) Free energy of partial nucleosome unwrapping for a system composed of the histone octamer plus 147-bp of DNA; WEChroM model. (d) Nucleosome AMH energy of the partial nucleosome unwrapping for the 147-bp WEChroM model. Example nucleosome configurations are shown for end-to-end distances of 76 Å (left), 142 Å (middle),221 Å (right)

We investigate the free energy landscape of DNA unwrapping from the nucleosome core, and its dependence on the parameters in our model. The reaction coordinate is chosen as the distance between the two free ends of the DNA, which mimics the nucleosome extension reported in the optical trap experiment(62). To explore the phase space of unwrapping more rapidly, we used Umbrella Sampling (US) and the Multiple Bennett Acceptance Ratio (MBAR) (63) to calculate the free energy as a function of end-to-end distance.

First we study a 223-bp system, with 147-bp DNA molecules bound to the histone core and 38-bp linker DNA particles on each side The same system was previously investigated using the 3-Site-per-nucleotide (3SPN) DNA model and the Atomic Interaction-based Coarse Grained (AICG) model of the nucleosome(32,64). We trained the parameters in *U*_*Nucleosome*_ to match the free energy computed with the more detailed 3SPN-AICG model.

After training, the free energy profile computed with WEChroM is in good agreement with the 3SPN-AICG (Fig. 4b right), despite the simplicity of the AMH energy function. Both models place the free energy minimum at around 80 Å and this free energy monotonically increases for the simulated range, 250 Å in the case of 3SPN-AICG and up to 350 Å in the current study. The extension energy in the full range is at the order of magnitude of a few *k*_*B*_*T*, matching experimental observations that nucleosomes can spontaneously unwrap and rewrap the outer layer of DNA(6).

With this trained set of parameters for WEChroM, we study the unwrapping of DNA from the nucleosome core particle, i.e., the histones octamer wrapped by 147-bp DNA. As before, the free energy profile is computed using Umbrella Sampling and MBAR. The results are displayed in Fig. 4c. Without the flexibility warranted by linker DNA, the free energy displays a two-staged non-monotonical increase as the end-to-end distance stretches to 220 Å, when the outer layer of DNA is completely dissociated from the histone core.

The AMH potential energy of the two-stage unwrapping process is shown in figure 4d. Consistently with the free energy, the nucleosome potential energy shows two sharp increases for the end-to-end distance between 80 and 100 Å a well as between 160 and 180 Å. An analysis of the structures associated with these end-to-end distances indicates that for each sharp increase, one end of the DNA is unbound from the histone core; snapshots of these conformations are displayed in figure 4d.

Altogether, our model suggests that the unwrapping of DNA from the nucleosome core particle is not a gradual – one base-pair at the time – process, instead, the molecular complex accumulates mechanical strain until one arm consisting of tens of base pairs suddenly breaks apart. The presence of molecular cracking in the DNA unwrapping process could be rationalized by considering the high stiffness of double-stranded DNA. On the other hand, because experiments typically require linker DNA to apply the external forces unwrapping the DNA, observing molecular cracking in the wet lab can be challenging, as we have shown that the presence of linker DNA masks this behavior. Overall, our results demonstrate that, beyond double-stranded DNA, the WEChroM model can effectively model the mechanics of single nucleosomes, opening the way to the study of the 10nm fiber and of genes.

## Discussion

The Widely Editable Chromatin Model (WEChroM) proposes an alternative approach to the modeling of chromatin and genes. The objective of this work is to reproduce the mechanical properties of the 10-nm fiber with a simple and computationally efficient mathematical model, with the compromise of relinquishing any attempt at reproducing the details of the chemical complexity of the system.

As shown, the model can reproduce the bending and twisting persistence lengths, and their temperature dependence. WEChroM is also able to reproduce the effects of positive and negative supercoiling, the formation of plectonemes and of structural defects that relax mechanical stress, and the asymmetric behavior with respect to positive or negative supercoiling, which was previously observed in experiments. Finally, we show that the proposed model is also suitable for reproducing the free energy of partial DNA unwrapping from nucleosomes. Indeed, these results demonstrate that the WEChroM model is a viable strategy to simulate the mechanics of 10-nm fiber, when willing to surrender chemical accuracy.

The structural ensembles of genes have been so far out of the reach of theoretical and computational investigations because genes are molecular complexes too big to be tackled with even the most efficient CG models and yet too strongly affected by heterogeneous molecular factors to be effectively modeled as a simple polymer. The WEChroM is an attempt to extend the reach of computational modeling with the alternative approach of AMH. Preliminary benchmarking (see Appendix S3) does indicate that the model is efficient enough to allow the study of genetic systems as big as tens of kilobases of chromatin, i.e., the size of mammalian genes.

Because of the extremely large number of combinations of nucleosome types, epigenetic modifications, linker lengths, and bound factors, modeling the chromatin fiber as a polymer composed of nucleosome monomers appears impractical. In fact, we are not likely to see the same exact monomer twice in a system composed of a few kilobases of DNA. WEChroM is also an attempt to solve this conundrum. The namesake characteristic of the Widely Editable Chromatin Model provides the needed flexibility for introducing a vast repertoire of structural memories of DNA binding proteins and histone variants and integrating the effect of these molecules at a larger scale. This characteristic of the AMH approach, which has not been yet exploited, opens the way to the study of structural ensembles of specific segments of chromatin, utilizing the vast amounts of omics data indicating the positioning and the identity of bound factors along DNA are already available for a variety of species, and human in particular.

## Materials and methods

### Implementation of WEChroM on OpenMM

In recent years, the OpenMM simulation toolkit was developed for high-performance and extensible molecular dynamics simulations(65). It provides a high-level application programming interface (API) for external programs like Python to utilize its functions. In the meanwhile, OpenMM makes the most efficient use of CPU and GPU hardware capabilities automatically with its lower layers.

WEChroM is implemented with a variety of custom force templates provided by the high-level API layer of OpenMM. For each term in the Hamiltonians, we select the force template that matches it the best. The code was written to support either naked DNA molecules or nucleosomes. Users can specify the hardware platform as an input parameter, and OpenMM compiles optimized code for the desired platform. Compatible platforms include single CPUs, multiple CPUs, and GPUs (with CUDA, OpenCL, and HIP support).

### Simulation Setup

The WEChroM software offers a function for coarse-graining atomic PDB files provided by the users into a WEChroM molecule. The atomic PDB files can be generated by X3DNA(47) or downloaded from the PDB Database. DNA particles and histone protein core particles (in the case of a nucleosome system) in the generated coarse-grained file will be automatically recognized by the software. This coarse-grained file will later be used as the initial structure of the simulation as well.

For naked DNA simulations, WEChroM is ready to run simulations using potential energy described in Section “Associative Memory Hamiltonian of DNA” after the initial preparation. The trajectories are saved in .dcd files. For trajectories reported in this work, each of them was simulated for 1 × 10^9^ steps, storing a frame every 5 × 10^3^ steps. Analyses are performed after the simulations by reading the trajectory files.

For nucleosome simulations, we employ umbrella sampling to sufficiently explore the configurational space. The DNA end-to-end distance serves as the reaction coordinate. The first or last base pair’s center of mass is used to designate the ends of the DNA, and the Euclidean distance between them is used to determine the end-to-end distance. Forty-one umbrella windows are used for the 223 bp system (nucleosome core particle with two arms), and 23 are used for the 147bp system (nucleosome core particle system). In each window, the end-to-end distance is restrained around a center distance using a harmonic biasing potential. Five replicas were used within each umbrella window, and each replica was simulated for 1 × 10^7^ steps storing a frame every 1 × 10^3^ steps. We then utilize the FastMBAR(63) to calculate the free energies and expectations based on the saved trajectories.

## Code Availability

WEChroM is available at https://github.com/VickyLu001/wechrom website. Usage tutorials for the package are available at https://wechrom.readthedocs.io/en/latest/ website.

## Acknowledgments

This research was supported by the Center for Theoretical Biological Physics sponsored by the NSF (Grants PHY-2019745 and PHY-2210291) and by the Welch Foundation (Grant C-1792). JNO is a CPRIT Scholar in Cancer Research sponsored by the Cancer Prevention and Research Institute of Texas. MDP is supported by the NIGMS of the National Institutes of Health under award number R35GM146852. The content is solely the responsibility of the authors and does not necessarily represent the official views of the National Institutes of Health. Finally, we thank AMD for providing the computational resources used in this research.

## Supporting information

**S1 Appendix. Simulation with Reduced Units**.

**S2 Appendix. Connectivity Interactions and Steric Interactions**.

**S3 Appendix. Comparison between WEChroM and Open3SPN2**.

**S4 Appendix. Calculations of the linking number, writhing number, and twisting number**.

**S1 Fig. Counting and the relative variance *σ* of associative memory terms**.

**S2 Fig. Model flexibility**.

**S3 Fig. Systems of different sizes are consistent in persistence length**.

**S1 Table. Parameters in *U***_***con***_ **and *U***_***νol***_.

**S2 Table. Multiple Sets of Parameters in *U***_***DH***_.

## References

1. Watson JD, Crick FHC. Molecular Structure of Nucleic Acids: A Structure for Deoxyribose Nucleic Acid. Nature. 1953 Apr;171(4356):737–8.

2. Franklin RE, Gosling RG. Molecular Configuration in Sodium Thymonucleate. Nature. 1953 Apr;171(4356):740–1.

3. Wilkins MHF, Stokes AR, Wilson HR. Molecular Structure of Nucleic Acids: Molecular Structure of Deoxypentose Nucleic Acids. Nature. 1953 Apr;171(4356):738–40.

4. Sun T, Minhas V, Korolev N, Mirzoev A, Lyubartsev AP, Nordenskiöld L. Bottom-Up Coarse-Grained Modeling of DNA. Front Mol Biosci. 2021;8.

5. Zhou K, Gaullier G, Luger K. Nucleosome structure and dynamics are coming of age. Nat Struct Mol Biol. 2019 Jan;26(1):3–13.

6. Tims HS, Gurunathan K, Levitus M, Widom J. Dynamics of Nucleosome Invasion by DNA Binding Proteins. J Mol Biol. 2011 Aug 12;411(2):430–48.

7. Lavery R, Zakrzewska K, Beveridge D, Bishop TC, Case DA, Cheatham T III, et al. A systematic molecular dynamics study of nearest-neighbor effects on base pair and base pair step conformations and fluctuations in B-DNA. Nucleic Acids Res. 2010 Jan 1;38(1):299–313.

8. Minhas V, Sun T, Mirzoev A, Korolev N, Lyubartsev AP, Nordenskiöld L. Modeling DNA Flexibility: Comparison of Force Fields from Atomistic to Multiscale Levels. J Phys Chem B. 2020 Jan 9;124(1):38–49.

9. Sun T, Mirzoev A, Minhas V, Korolev N, Lyubartsev AP, Nordenskiöld L. A multiscale analysis of DNA phase separation: from atomistic to mesoscale level. Nucleic Acids Res. 2019 Jun 20;47(11):5550–62.

10. Savelyev A, Papoian GA. Molecular Renormalization Group Coarse-Graining of Polymer Chains: Application to Double-Stranded DNA. Biophys J. 2009 May 20;96(10):4044–52.

11. Kovaleva NA, Koroleva (Kikot) IP, Mazo MA, Zubova EA. The “sugar” coarse-grained DNA model. J Mol Model. 2017 Feb 9;23(2):66.

12. Naômé A, Laaksonen A, Vercauteren DP. A Coarse-Grained Simulation Study of the Structures, Energetics, and Dynamics of Linear and Circular DNA with Its Ions. J Chem Theory Comput. 2015 Jun 9;11(6):2813–26.

13. Maffeo C, Ngo TTM, Ha T, Aksimentiev A. A Coarse-Grained Model of Unstructured Single-Stranded DNA Derived from Atomistic Simulation and Single-Molecule Experiment. J Chem Theory Comput. 2014 Aug 12;10(8):2891–6.

14. Ouldridge TE, Louis AA, Doye JPK. Structural, mechanical, and thermodynamic properties of a coarse-grained DNA model. J Chem Phys. 2011 Feb 28;134(8):085101.

15. Snodin BEK, Randisi F, Mosayebi M, Šulc P, Schreck JS, Romano F, et al. Introducing improved structural properties and salt dependence into a coarse-grained model of DNA. J Chem Phys. 2015 Jun 21;142(23):234901.

16. Knotts TA, Rathore N, Schwartz DC, de Pablo JJ. A coarse grain model for DNA. J Chem Phys. 2007 Feb 28;126(8):084901.

17. Sambriski EJ, Schwartz DC, de Pablo JJ. A Mesoscale Model of DNA and Its Renaturation. Biophys J. 2009 Mar 4;96(5):1675–90.

18. Hinckley DM, Freeman GS, Whitmer JK, de Pablo JJ. An experimentally-informed coarse-grained 3-site-per-nucleotide model of DNA: Structure, thermodynamics, and dynamics of hybridization. J Chem Phys. 2013 Oct 14;139(14):144903.

19. Freeman GS, Hinckley DM, Lequieu JP, Whitmer JK, de Pablo JJ. Coarse-grained modeling of DNA curvature. J Chem Phys. 2014 Oct 28;141(16):165103.

20. Markegard CB, Fu IW, Reddy KA, Nguyen HD. Coarse-Grained Simulation Study of Sequence Effects on DNA Hybridization in a Concentrated Environment. J Phys Chem B. 2015 Feb 5;119(5):1823–34.

21. Darré L, Machado MR, Brandner AF, González HC, Ferreira S, Pantano S. SIRAH: A Structurally Unbiased Coarse-Grained Force Field for Proteins with Aqueous Solvation and Long-Range Electrostatics. J Chem Theory Comput. 2015 Feb 10;11(2):723–39.

22. Cragnolini T, Derreumaux P, Pasquali S. Coarse-Grained Simulations of RNA and DNA Duplexes. J Phys Chem B. 2013 Jul 11;117(27):8047–60.

23. Dans PD, Walther J, Gómez H, Orozco M. Multiscale simulation of DNA. Curr Opin Struct Biol. 2016 Apr 1;37:29–45.

24. He Y, Maciejczyk M, Ołdziej S, Scheraga HA, Liwo A. Mean-Field Interactions between Nucleic-Acid-Base Dipoles can Drive the Formation of a Double Helix. Phys Rev Lett. 2013 Feb 28;110(9):098101.

25. Perišić O, Collepardo-Guevara R, Schlick T. Modeling Studies of Chromatin Fiber Structure as a Function of DNA Linker Length. J Mol Biol. 2010 Nov 12;403(5):777–802.

26. Ozer G, Luque A, Schlick T. The chromatin fiber: multiscale problems and approaches. Curr Opin Struct Biol. 2015 Apr 1;31:124–39.

27. Arya G, Schlick T. A Tale of Tails: How Histone Tails Mediate Chromatin Compaction in Different Salt and Linker Histone Environments. J Phys Chem A. 2009 Apr 23;113(16):4045–59.

28. Perišić O, Schlick T. Dependence of the Linker Histone and Chromatin Condensation on the Nucleosome Environment. J Phys Chem B. 2017 Aug 24;121(33):7823–32.

29. Bascom GD, Kim T, Schlick T. Kilobase Pair Chromatin Fiber Contacts Promoted by Living-System-Like DNA Linker Length Distributions and Nucleosome Depletion. J Phys Chem B. 2017 Apr 20;121(15):3882–94.

30. Bascom GD, Sanbonmatsu KY, Schlick T. Mesoscale Modeling Reveals Hierarchical Looping of Chromatin Fibers Near Gene Regulatory Elements. J Phys Chem B. 2016 Aug 25;120(33):8642–53.

31. Bascom GD, Myers CG, Schlick T. Mesoscale modeling reveals formation of an epigenetically driven HOXC gene hub. Proc Natl Acad Sci. 2019 Mar 12;116(11):4955–62.

32. Lequieu J, Córdoba A, Moller J, de Pablo JJ. 1CPN: A coarse-grained multi-scale model of chromatin. J Chem Phys. 2019 Jun 7;150(21):215102.

33. Davtyan A, Schafer NP, Zheng W, Clementi C, Wolynes PG, Papoian GA. AWSEM-MD: Protein Structure Prediction Using Coarse-Grained Physical Potentials and Bioinformatically Based Local Structure Biasing. J Phys Chem B. 2012 Jul 26;116(29):8494–503.

34. Dekker J, Rippe K, Dekker M, Kleckner N. Capturing Chromosome Conformation. Science. 2002 Feb 15;295(5558):1306–11.

35. Zhao Z, Tavoosidana G, Sjölinder M, Göndör A, Mariano P, Wang S, et al. Circular chromosome conformation capture (4C) uncovers extensive networks of epigenetically regulated intra- and interchromosomal interactions. Nat Genet. 2006 Nov;38(11):1341–7.

36. Lieberman-Aiden E, van Berkum NL, Williams L, Imakaev M, Ragoczy T, Telling A, et al. Comprehensive Mapping of Long-Range Interactions Reveals Folding Principles of the Human Genome. Science. 2009 Oct 9;326(5950):289–93.

37. Quinodoz SA, Ollikainen N, Tabak B, Palla A, Schmidt JM, Detmar E, et al. Higher-Order Inter-chromosomal Hubs Shape 3D Genome Organization in the Nucleus. Cell. 2018 Jul 26;174(3):744-757.e24.

38. Beliveau BJ, Joyce EF, Apostolopoulos N, Yilmaz F, Fonseka CY, McCole RB, et al. Versatile design and synthesis platform for visualizing genomes with Oligopaint FISH probes. Proc Natl Acad Sci. 2012 Dec 26;109(52):21301–6.

39. Boettiger AN, Bintu B, Moffitt JR, Wang S, Beliveau BJ, Fudenberg G, et al. Super-resolution imaging reveals distinct chromatin folding for different epigenetic states. Nature. 2016 Jan;529(7586):418–22.

40. Fabre PJ, Benke A, Joye E, Nguyen Huynh TH, Manley S, Duboule D. Nanoscale spatial organization of the HoxD gene cluster in distinct transcriptional states. Proc Natl Acad Sci. 2015 Nov 10;112(45):13964–9.

41. Mateo LJ, Murphy SE, Hafner A, Cinquini IS, Walker CA, Boettiger AN. Visualizing DNA folding and RNA in embryos at single-cell resolution. Nature. 2019 Apr;568(7750):49–54.

42. Di Pierro M, Zhang B, Aiden EL, Wolynes PG, Onuchic JN. Transferable model for chromosome architecture. Proc Natl Acad Sci. 2016 Oct 25;113(43):12168–73.

43. Oliveira Junior AB, Contessoto VG, Mello MF, Onuchic JN. A Scalable Computational Approach for Simulating Complexes of Multiple Chromosomes. J Mol Biol. 2021 Mar 19;433(6):166700.

44. MacPherson Q, Beltran B, Spakowitz AJ. Bottom–up modeling of chromatin segregation due to epigenetic modifications. Proc Natl Acad Sci. 2018 Dec 11;115(50):12739–44.

45. Friedrichs MS, Wolynes PG. Toward Protein Tertiary Structure Recognition by Means of Associative Memory Hamiltonians. Science. 1989 Oct 20;246(4928):371–3.

46. Friedrichs MS, Wolynes PG. Molecular dynamics of associative memory hamiltonians for protein tertiary structure recognition. Tetrahedron Comput Methodol. 1990 Jan 1;3(3):175–90.

47. Li S, Olson WK, Lu XJ. Web 3DNA 2.0 for the analysis, visualization, and modeling of 3D nucleic acid structures. Nucleic Acids Res. 2019 Jul 2;47(W1):W26–34.

48. Gray HB, Hearst JE. Flexibility of native DNA from the sedimentation behavior as a function of molecular weight and temperature. J Mol Biol. 1968 Jul 14;35(1):111–29.

49. Geggier S, Kotlyar A, Vologodskii A. Temperature dependence of DNA persistence length. Nucleic Acids Res. 2011 Mar 1;39(4):1419–26.

50. Baumann CG, Smith SB, Bloomfield VA, Bustamante C. Ionic effects on the elasticity of single DNA molecules. Proc Natl Acad Sci. 1997 Jun 10;94(12):6185–90.

51. Moroz JD, Nelson P. Torsional directed walks, entropic elasticity, and DNA twist stiffness. Proc Natl Acad Sci. 1997 Dec 23;94(26):14418–22.

52. Noy A, Golestanian R. Length Scale Dependence of DNA Mechanical Properties. Phys Rev Lett. 2012 Nov 30;109(22):228101.

53. Gao X, Hong Y, Ye F, Inman JT, Wang MD. Torsional Stiffness of Extended and Plectonemic DNA. Phys Rev Lett. 2021 Jul 7;127(2):028101.

54. Roth E, Glick Azaria A, Girshevitz O, Bitler A, Garini Y. Measuring the Conformation and Persistence Length of Single-Stranded DNA Using a DNA Origami Structure. Nano Lett. 2018 Nov 14;18(11):6703–9.

55. Fogg JM, Randall GL, Pettitt BM, Sumners DWL, Harris SA, Zechiedrich L. Bullied no more: when and how DNA shoves proteins around. Q Rev Biophys. 2012 Aug;45(3):257–99.

56. Seidel R, Dekker C. Single-molecule studies of nucleic acid motors. Curr Opin Struct Biol. 2007 Feb 1;17(1):80–6.

57. Harada Y, Ohara O, Takatsuki A, Itoh H, Shimamoto N, Kinosita K. Direct observation of DNA rotation during transcription by Escherichia coli RNA polymerase. Nature. 2001 Jan;409(6816):113–5.

58. Irobalieva RN, Fogg JM, Catanese DJ, Sutthibutpong T, Chen M, Barker AK, et al. Structural diversity of supercoiled DNA. Nat Commun. 2015 Oct 12;6(1):8440.

59. Bettotti P, Visone V, Lunelli L, Perugino G, Ciaramella M, Valenti A. Structure and Properties of DNA Molecules Over The Full Range of Biologically Relevant Supercoiling States. Sci Rep. 2018 Apr 18;8(1):6163.

60. Meng H, Bosman J, van der Heijden T, van Noort J. Coexistence of Twisted, Plectonemic, and Melted DNA in Small Topological Domains. Biophys J. 2014 Mar 4;106(5):1174–81.

61. Li D, Lv B, wang Q, Liu Y, Zhuge Q. Direct observation of positive supercoils introduced by reverse gyrase through atomic force microscopy. Bioorg Med Chem Lett. 2017 Sep 1;27(17):4086–90.

62. Mihardja S, Spakowitz AJ, Zhang Y, Bustamante C. Effect of force on mononucleosomal dynamics. Proc Natl Acad Sci. 2006 Oct 24;103(43):15871–6.

63. Ding X, Vilseck JZ, Brooks CL. Fast Solver for Large Scale Multistate Bennett Acceptance Ratio Equations. J Chem Theory Comput. 2019 Feb 12;15(2):799–802.

64. Lequieu J, Córdoba A, Schwartz DC, de Pablo JJ. Tension-Dependent Free Energies of Nucleosome Unwrapping. ACS Cent Sci. 2016 Sep 28;2(9):660–6.

65. Eastman P, Swails J, Chodera JD, McGibbon RT, Zhao Y, Beauchamp KA, et al. OpenMM 7: Rapid development of high performance algorithms for molecular dynamics. PLOS Comput Biol. 2017 Jul 26;13(7):e1005659.

