## Supporting Information for "An Associative Memory Hamiltonian Model for DNA and Nucleosomes"

**S1 Appendix. Simulation with Reduced Units.** In our simulations, the length in the WEChroM system is expressed in relative units, where the equilibrium distance between two adjunct particles on the same strand is 2.0 (around 0.482 nm in standard units). The energy is expressed in the unit of $k_B T$, where the temperature ($T$) of the simulation is fixed and assumed to be room temperature, and we tune the energy terms relatively.

**S2 Appendix. Connectivity Interactions and Steric Interactions.**

The two basic polymer parts of the potential energy are the connected-chain terms $U_{con}$ and steric terms $U_{vol}$. The connected-chain term $U_{con}$ is a harmonic spring between the nearest particles on the same strand given by

$$U_{con} = \frac{1}{2} k_{con} \left( r_{i,i+1} - r_0 \right)^2 \tag{1}$$

where $r_{i,i+1}$ is the distance between the nearest particles on the same strand, $k_{con}$ is the harmonic force constant, and $r_0$ is the equilibrium position distance between two particles and is set to 2.0 in the reduced unit system. It plays the role of forming backbone covalent bonds between the two neighbor nucleotides which should not break under room temperature.

The steric energy $U_{vol}$ is short-range repulsive energy between any pair of particles to prevent them from overlapping with each other, excepting the pair of the two nearest particles on the same strand, which are already constrained by the harmonic springs of $U_{con}$. It is given by

$$U_{vol} = k_{vol}(r_{ij} - r^*)^2 H(r^* - r_{ij})$$

(2)

where $k_{vol}$ is the harmonic force constant, $r_{ij}$ is the distance between particle *i* and particle *j* to which the energy is applied, and $r^*$ is approximately twice the effective radius of the particle and is set to be 2.07, *H(x)=1* for *x>0*, *H(x)=0* otherwise.

These two energies form the "backbone" of our DNA system, and with these terms the particle chain can maintain itself with enough rotation freedom to form a certain structure given additional Hamiltonian terms. It is clear that $U_{con}$ should be much larger than the thermal energy $k_BT$ when $\Delta r = (r - r_0)$ is comparable to $r_0$, because in practice the DNA single chain can hardly be broken at room temperature due to the covalent bond connecting the nucleotides. In our model, we set the $k_{con}$ on such a scale that when $\Delta r \sim 0.01 r_0$, $U_{con}$ is comparable to $k_BT$. Accordingly, our relative bond length variance between the nearest particles is $\sim 0.01$, consistent with previous reports and simulations(1). Detailed parameters are provided in S1 Table.

**S3 Appendix. Comparison between WEChroM and Open3SPN2.**

As a more coarse-grained model, our model gives better computing efficiency than that of 3SPN.2(1). We performed simulations of 250-bp naked DNA supercoiling for 24 hours based on the same environment (NVIDIA Tesla V100, CUDA version 10.1, OpenMM version 7.4) using these two models. Our model executed 1.02e9 steps while the 3SPN model executed 5.4e7 steps. It is clear that our model is nearly 20 times faster than 3SPN in terms of execution steps.

**S4 Appendix. Calculations of the linking number, writhing number, and twisting number.**

The linking number describes the number of times that one curve winds around the other. Mathematically, the linking number of two closed curves $C_1$ and $C_2$ is

$$Lk = \frac{1}{4\pi}\int_{C1}\int_{C2} d\Omega(\boldsymbol{r}_1, \boldsymbol{r}_2)$$

( 3 )

where $\boldsymbol{r}_1$ and $\boldsymbol{r}_2$ are two arbitrary points passing along the curves $C_1$ and $C_2$ and the solid angle $d\Omega(\boldsymbol{r}_1, \boldsymbol{r}_2)$ is defined by

$$d\Omega(\mathbf{r_1}, \mathbf{r_2}) = \frac{(d\boldsymbol{r}_2 \times d\boldsymbol{r}_1)\boldsymbol{r}_{12}}{r_{12}^3}$$

( 4 )

where $d\boldsymbol{r}_1$ and $d\boldsymbol{r}_2$ are two infinitesimal vectors originating from $\boldsymbol{r}_1$ and $\boldsymbol{r}_2$, the vector $\boldsymbol{r}_{12} = \boldsymbol{r}_2 - \boldsymbol{r}_1$, and $r_{12} = |\boldsymbol{r}_{12}|$. The writhe of a curve C measures the number of times that one curve winds around itself, and the mathematical formula is

$$Wr = \frac{1}{4\pi}\int_{C}\int_{C} d\Omega(\boldsymbol{r}_1, \boldsymbol{r}_2)$$

( 5 )

where $d\Omega(\boldsymbol{r}_1, \boldsymbol{r}_2)$ is defined by Eq. 12 and $d\boldsymbol{r}_1$ and $d\boldsymbol{r}_2$ pass along the curve $C$.

The Gauss double integral (Eq. 11) along a DNA polygon of $N$ segments can be expressed as the double sum,

$$Lk = \sum_{i=1}^{N}\sum_{j=1}^{N}\frac{\Omega_{ij}}{4\pi}$$

( 6 )

where $i$ and $j$ are the DNA particles on the two strands respectively. The writhing number in the DNA system is

$$Wr = 2\sum_{i=1}^{N}\sum_{j>i}^{N}\frac{\Omega_{ij}}{4\pi} \quad (7)$$

where *i* and *j* pass along the centerline of the DNA, defined by each basepair's center of mass. To calculate the solid angle $\Omega_{ij}$ in the DNA system, we use method 1a provided by Konstantin Klenin and Jörg Langowski(2). The twisting number measures the number of rotations the DNA polymer possesses and can be derived by $Tw = Lk - Wr$.

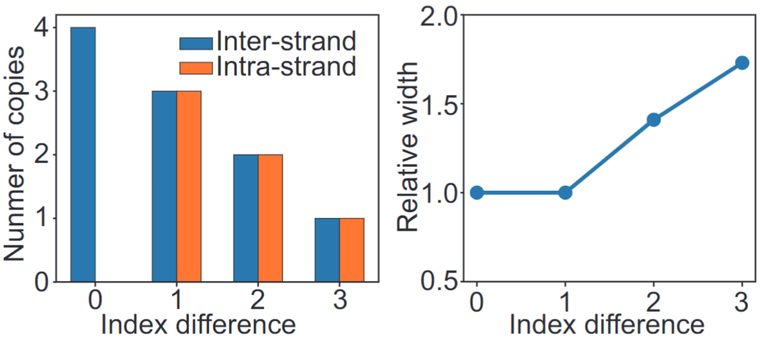


**S1 Fig. Counting and the relative variance $\sigma$ of associative memory terms.** (a) Due to the nature of counting, the number of associative memory bonds applied to a pair of particles depends on the index difference. (b) We expand the relative variance $\sigma$ of the bonds depending on the index difference in the trend of $\sqrt{index\ difference}$

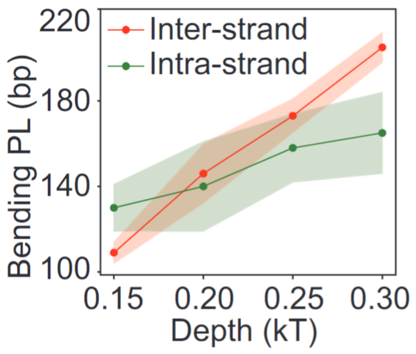


**S2 Fig. Model flexibility.** The figure shows how bending persistence lengths depend on the two types of energy scaling factor $\lambda$. The solid lines are persistence length for inter-strand (red) $\lambda$ and intra-strand factor $\lambda$ (green) and the shaded area are standard error. The variance $\sigma$ is fixed at 0.15. In each line, only one $\lambda$ is varied and the other is fixed.

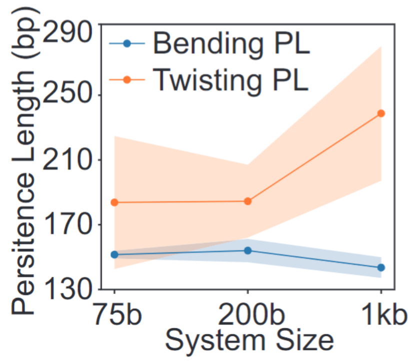


**S3 Fig. Systems of different sizes are consistent in persistence length.** The figure shows how twisting and bending persistence lengths depend on the size of the WEChroM system. The solid lines are persistence lengths for bending (blue) and twisting (orange) on different system sizes and the shaded area is the standard error.

| Parameter (unit) | Value |
|---|---|
| $k_{con}$ ($k_B T$) | 3000 |

| | |
|---|---|
| $r_0$ | 2.0 |
| $k_{vol}$ ($k_BT$) | 5856 |
| $r*$ | 2.07 |

**S1 Table. Parameters in $U_{con}$ and $U_{vol}$.** We used the reduced units, where the equilibrium distance between two adjunct particles on the same strand is 2.0 (around 0.482 nm in standard units). The energy is expressed in the unit of $k_BT$, where the temperature ($T$) of the simulation is fixed and assumed to be room temperature, and we tune the energy terms relatively. The parameters used in $U_{con}$ and $U_{vol}$ are summarized in the table.

| λ ($k_BT$) | σ (r.u) | $L_{bp}$ (bp) | $L_{tp}$ (bp) |
|---|---|---|---|
| 0.15 | 0.05 | *144* | *188* |
| 0.15 | 0.1 | 153 | 172 |
| 0.2 | 0.15 | 140 | 191 |
| 0.25 | 0.17 | 154 | 196 |
| 0.3 | 0.2 | 151 | 219 |

**S2 Table. Multiple Sets of Parameters in $U_{DH}$.** These sets of parameters all give a bending persistence length $L_{bp}$ ~ 150 bp.

### References


1. Hinckley DM, Freeman GS, Whitmer JK, de Pablo JJ. An experimentally-informed coarse-grained 3-site-per-nucleotide model of DNA: Structure, thermodynamics, and dynamics of hybridization. J Chem Phys. 2013 Oct 14;139(14):144903.

2. Klenin K, Langowski J. Computation of writhe in modeling of supercoiled DNA. Biopolymers. 2000;54(5):307–17.
